## Supplementary figures for "Somatodendritic orientation determines tDCS-induced neuromodulation of Purkinje cell activity in awake mice"

### Supplementary figure legends

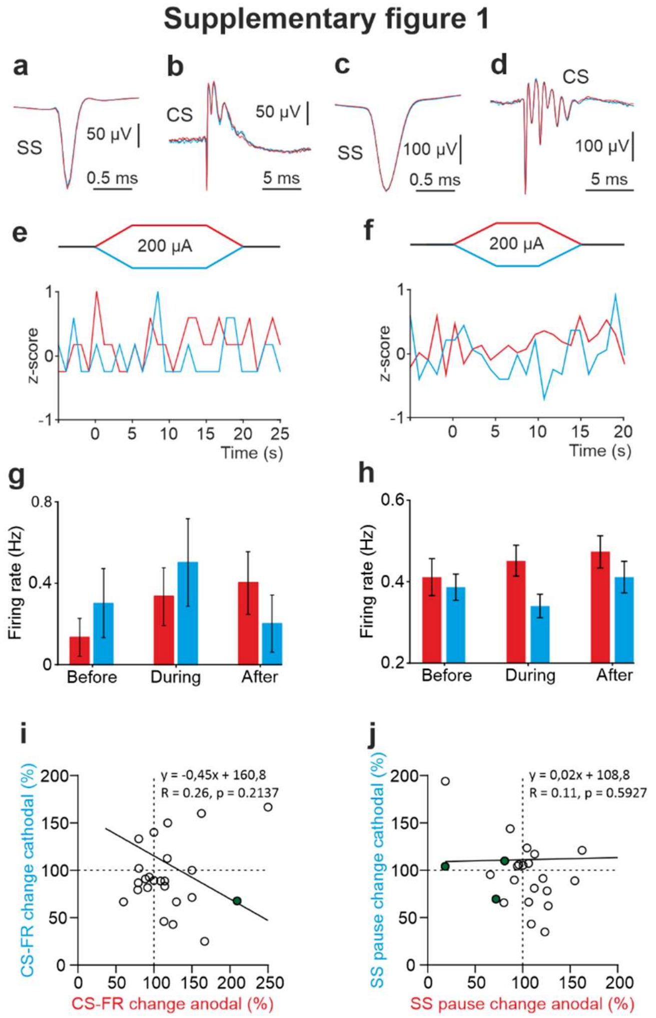


**Supplementary fig. 1 tDCS does not modulate PC waveform or complex spikes in the awake mouse. a-d** Superimposed averaged SS (a, c) and CS (b, d) waveforms under control (black), anodal (red) and cathodal (blue) tDCS. **e,f** Z-score-transformed average PSTH (bin size: 1 s) of CS activity before, during and after anodal (red trace) or cathodal (blue trace) tDCS corresponding to PC showed in fig. 2a and fig. 2b in the main text, respectively. **g,h** Statistical comparison of CS firing rate between 5 s windows before, during, and after tDCS (**g**: Anodal: Friedman, $\chi2$ (3, 30) = 2.790, N = 11, p = 0.425; Cathodal: Friedman, $\chi2$ (3, 21) = 2.739, N = 8, p = 0.434; **h**: Anodal: Friedman, $\chi2$ (2, 108) = 2.045, N = 55, p = 0.359; Cathodal: RM-ANOVA, F (2, 110) = 1.706, N = 56, p = 0.426). Error bars represent SEM. **p < 0.01; ***p < 0.001. **i-j** Modulation of CS (i) firing rate and SS silence after a CS (j) of individual neurons (circles) during anodal (red) and cathodal (blue) tDCS. Filled circles represent statistically significant modulation during tDCS (n= 24, RM-ANOVA or Friedman tests, p < 0.05).

#### Supplementary figure 2

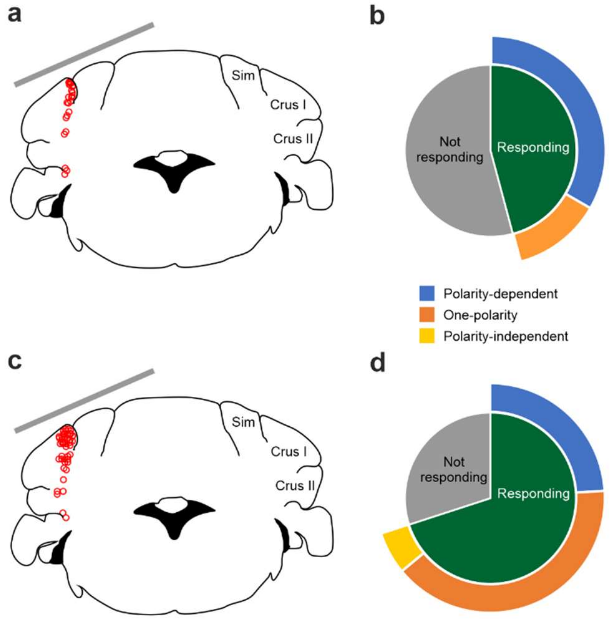


**Supplementary fig. 2 Summary of firing rate modulation for all recorded PCs and non-PCs during tDCS over crus I-II cerebellar region. a** Schematic representation of the recording sites of PCs and active electrode location (gray bar) during tDCS. **b** 45.8% of the recorded PC significantly modified their SS firing rate during tDCS (n = 11 PCs; green bar). The observed modulation consisted of a heterogeneous effect, with 12.5% of neurons significantly responding in a polarity-dependent manner to anodal and cathodal tDCS, with opposing firing rate changes (n = 3 PCs; blue bar) and 33.3% of the neurons significantly responding only to one tDCS polarity (n = 8 PCs; orange bar). **c** Schematic representation of the recording sites of non-PCs and active electrode location (gray bar) during tDCS. **d** 70% of recorded non-PC neurons significantly modified their firing rate during simultaneous tDCS (n = 35 non-PCs, green bar). Similar to PC, the observed modulation consisted of a heterogeneous effect, with 24% of neurons significantly responding in a polarity-dependent manner to both anodal and cathodal tDCS with opposing firing changes (n = 12 non-PCs; blue bar), 40% of the neurons significantly responding only to one tDCS polarity (n = 20 non-PCs; orange bar) and 6% significantly responding homogeneously to both tDCS polarities (n = 3 PCs; yellow bar i). (RM-ANOVA or Friedman tests, $p < 0.05$).

#### Supplementary figure 3

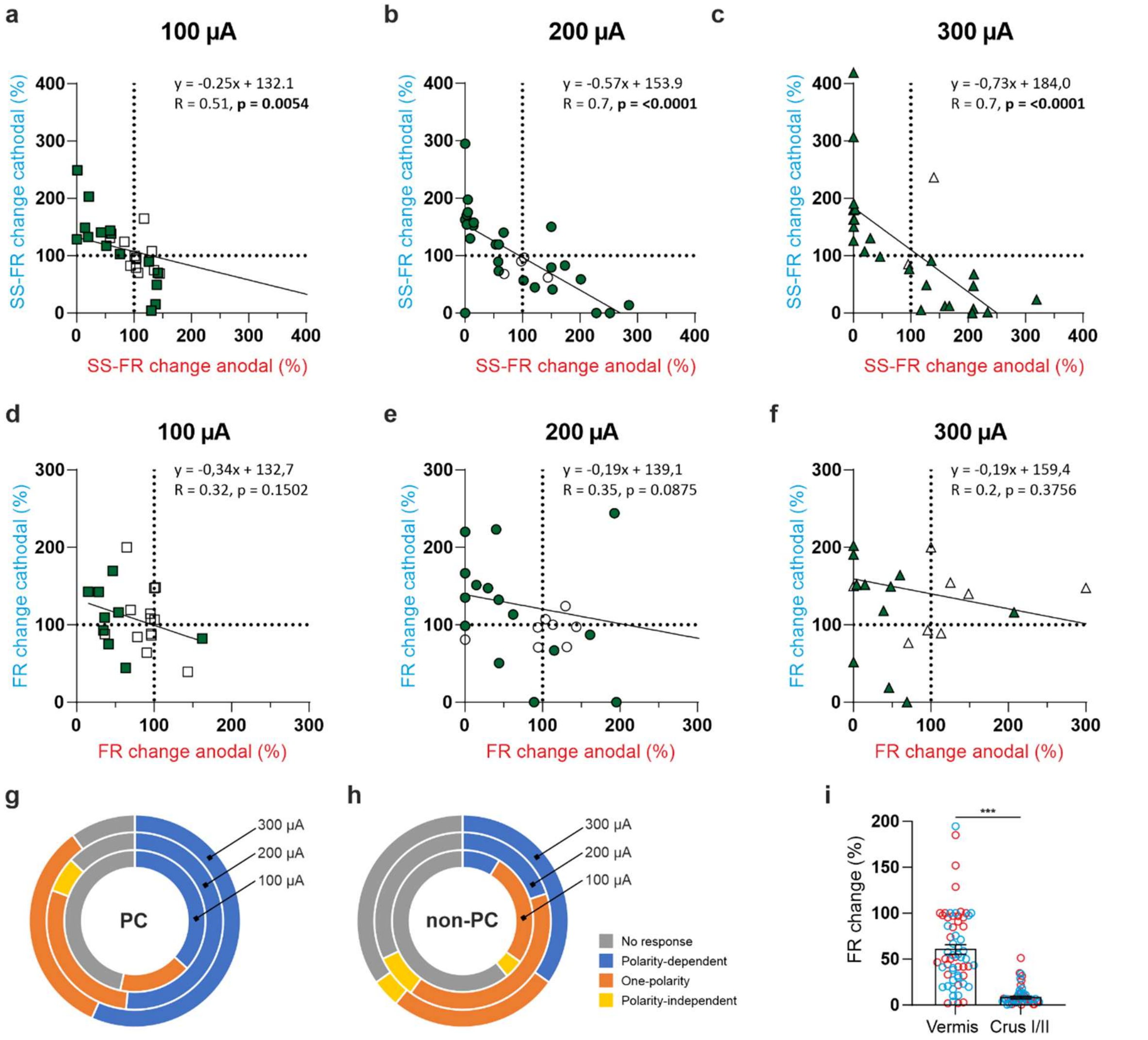


**Supplementary fig. 3 tDCS modulation of PC and non-PC activity at different intensities in anesthetized mice. a-c** Modulation of SS firing rate of the same individual PC during anodal (red) and cathodal (blue) tDCS over cerebellar vermis at different intensities (±100 µA, ±200 µA, and ±300 µA). Filled circles represent statistically significant modulation during tDCS (n = 31 PCs, RM-ANOVA or Friedman tests, $p < 0.05$). **d-f** Modulation of SS firing rate of the same individual non-PC during anodal (red) and cathodal (blue) tDCS over cerebellar vermis at different intensities (±100 µA, ±200 µA, and ±300 µA). Filled circles represent statistically significant modulation during tDCS (n = 25 non-PCs, RM-ANOVA or Friedman tests, $p < 0.05$). **g,h** Summary of SS firing rate modulation for all recorded PCs (g) and non-PCs (h) during tDCS over vermis cerebellar region. Neurons are grouped according to their modulatory response to tDCS. **i** Absolute firing rate modulation during anodal (red circles) and cathodal (blue circles) tDCS over Vermis and Crus I/II. Error bars represent SEM. ***$p < 0.001$ (Mann-Whitney test). **b** and **e** figures correspond to Fig. 4c and 4d respectively.

#### Supplementary figure 4

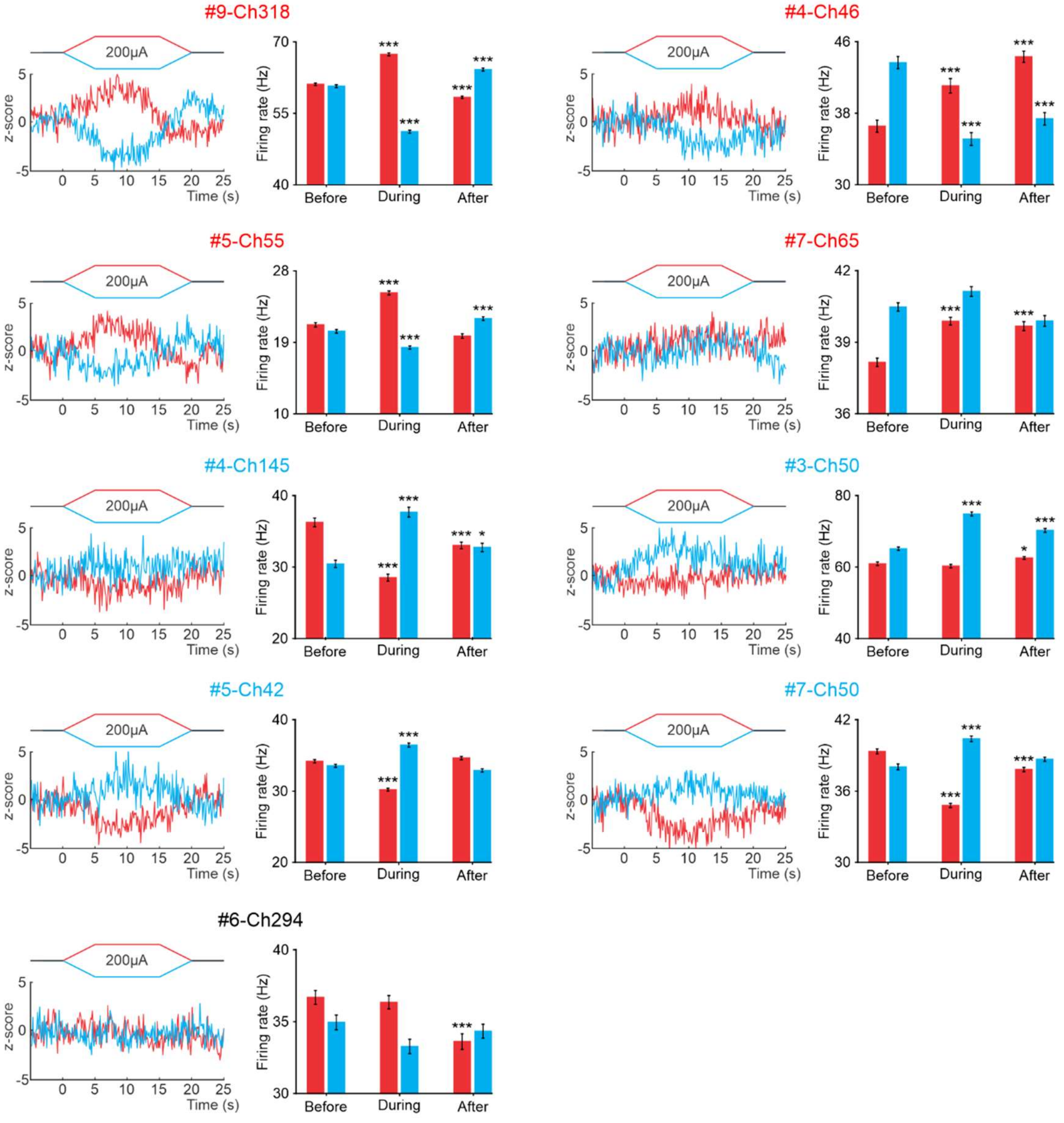


**Supplementary fig. 4 tDCS modulation of PCs at different PC layers in the awake mice.** The figure displays Z-score-transformed average PSTH (bin size: 0.1 s for SS) of the spontaneous SS recorded before, during and after anodal and cathodal tDCS pulses for each the recorded PCs included in the figure 6. Additionally, a statistical comparison of the SS firing rates during 5-second windows before, during, and after tDCS is presented for each recorded PC **(#9-Ch318**: Anodal: RM-ANOVA, F (2, 147) = 302.050, N = 50, p < 0.001; Cathodal: RM-ANOVA, F (2, 147) = 495.481, N = 50, p < 0.001; **#4-Ch46**: Anodal: RM-ANOVA, F (2, 147) = 30.325, N = 50, p < 0.001; Cathodal: RM-ANOVA, F (2, 147) = 40.236, N = 50, p < 0.001; **#5-Ch55**: Anodal: Friedman, $\chi^2$ (2, 98) = 67.647, N = 50, p < 0.001; Cathodal: Friedman, $\chi^2$ (2, 98) = 54.576, N = 50, p < 0.001; **#7-Ch65**: Anodal: RM-ANOVA, F (2, 147) = 28.562, N = 50, p < 0.001; Cathodal: RM-ANOVA, F (2, 147) = 9.282, N = 50, p < 0.001; **#4-Ch145**: Anodal: RM-ANOVA, F (2, 147) = 55.200, N = 50, p < 0.001;

Cathodal: RM-ANOVA, $F (2, 147) = 38.199$, $N = 50$, $p < 0.001$; **#3-Ch50**: Anodal: RM-ANOVA, $F (2, 147) = 6.257$, $N = 50$, $p = 0.003$; Cathodal: RM-ANOVA, $F (2, 147) = 86.379$, $N = 50$, $p < 0.001$; **#5-Ch42**: Anodal: RM-ANOVA, $F (2, 147) = 123.947$, $N = 50$, $p < 0.001$; Cathodal: RM-ANOVA, $F (2, 147) = 63.076$, $N = 50$, $p < 0.001$; **#7-Ch50**: Anodal: Friedman, $\chi^2 (2, 98) = 73.156$, $N = 50$, $p < 0.001$; Cathodal: Friedman, $\chi^2 (2, 98) = 32.827$, $N = 50$, $p < 0.001$; **#6-Ch294**: Anodal: RM-ANOVA, $F (2, 147) = 11.444$, $N = 50$, $p < 0.001$ Cathodal: Friedman, $\chi^2 (2, 98) = 4.135$, $N = 50$, $p = 0.127$). The error bars represent SEM. Significance levels are denoted as follows: *$p < 0.05$; ***$p < 0.001$.
